# Highlighting the complexity of pathogenesis: the host microbiota impacts disease development on apple fruit and is a cornerstone for its biocontrol

**DOI:** 10.1101/2024.08.21.608933

**Authors:** Abdoul Razack Sare, M. Haissam Jijakli, Sébastien Massart

## Abstract

Apple fruit is the most produced temperate fruit with a trade value estimated at 7.5 billion $ and is usually stored up to one year after harvest. Postharvest pathogens often compromise storage, responsible for up to 55% of fruit losses, depending on the country and fruit. They are also a source of mycotoxin contamination. A sustainable way to control that pathogen is using beneficial microorganisms called biocontrol agents (BCA). Despite promising efficacy in laboratory conditions, BCA’s efficacy is variable and often reduced once applied at a large scale through either orchard or postharvest treatment. We hypothesized that the epiphytic microbiota plays a role in the variability of BCA efficiency (*Pichia anomala*, strain K) and postharvest disease development due to *Botrytis cinerea* on apples.

A diverse set of 18 epiphytic microbial communities were harvested from apple carposphere and bio-banked. The analysis of their bacterial and fungal taxonomic composition and carbon metabolic footprint confirmed that contrasted microbiotas were harvested. Their impact on *B. cinerea* disease development was evaluated through a standardized *in vivo* bioassay. The reduction of *B. cinerea* rot development ranged from 20% to 80% when the microbiotas were applied alone. In addition, three microbiotas enhanced the biological control efficiency of strain K (up to +100%, whatever the tested microbiota concentrations) while others limited its action (down to -27%). A co-clustering analysis of biocontrol efficacy with carbon profiles or taxonomic composition was carried out. It identified promising molecules whose high metabolization was associated with high biocontrol by the microbiota and taxa with higher abundance in microbiota limiting *B. cinerea* rot development. Putative beneficial taxa were isolated from the most efficient microbiota. *In vivo* bioassays confirmed the efficacy of two molecules and two strains belonging to species never mentioned for their biocontrol properties against plant disease.

This study demonstrated that natural epiphytic microbiota significantly influences postharvest disease development in apples and cause a variability in biocontrol efficacy. By mining the generated data, our approach identified promising molecules and taxa that enhance biocontrol, offering new insights for sustainable postharvest pathogen management.

## Introduction

In plant pathology, the traditional “disease triangle” model explains pathogenesis by considering three key factors: a susceptible host, a virulent pathogen, and conducive environmental conditions. However, recent research suggests that this model is insufficient on its own, as plant health is also heavily influenced by its associated microbial communities, both epiphytic (living on the plant’s surface) and endophytic (living inside the plant). These plant-associated microbiota interact with the plant and the pathogen while being influenced by the environment (1). The contribution of plant-associated microbiota to disease development has led to the concept of pathobiome that integrates the plant-pathogen interactions within their biotic environment and the environmental conditions (2). Numerous studies have shown that pathogen development can alter plant-associated microbiota, with variations depending on the plant organ affected (3). These changes have been exploited to isolate beneficial microbial strains and design synthetic communities to enhance disease resistance (4, 5).

The impact of the residing microbiota on disease development has been already postulated (6) but very rarely studied, except for example, an observational study suggested a crucial role of rhizosphere protists in plant health (7). On the other hand, in several publications, modifying the environment by applying carbon or nitrogen sources of mineral or organic origin reshaped the initial microbiota, impacting pathogen or disease development (8, 9).

The postharvest losses threaten food security and sustainability, with up to 55% food losses (10). They remained stable during the past decades despite developing and applying chemical, physical, and biological treatments. Apple, the most-produced temperate fruit, hosts an epiphytic microbiota whose composition has already been characterized in contrasted agricultural conditions worldwide (11). Genera of postharvest pathogens such as *Botrytis*, *Monilinia*, *Neofabraea*, and *Penicillium* have been frequently observed, and their detection frequency tended to rise during storage (12), even without disease development. So, understanding the interaction between pathogens and the epiphytic microbiota is promising for developing new biocontrol solutions.

Many biocontrol agents (BCA) were isolated from apple epiphytic microbiota and developed into microbial-based biopesticides. However, despite decades of research, their efficacy is unstable and reduced compared to synthetic pesticides (13). Beyond well-studied abiotic conditions, this instability might also be due to complex ecological interactions with the resident microbiota. So, shifting from the disease triangle to the pathobiome paradigm is a promising axis to improve the control of diseases in terms of efficacy and reliability, also postharvest (14, 15).

The impact of BCA on apple fruit (16) or flower (17) epiphytic microbiota has already been studied. Nevertheless, apart from hypotheses, little information is available on the impact of the epiphytic microbiota on the efficacy of beneficial microorganisms like BCAs, whatever the plant species. This information is essential for understanding the ecological fitness of BCA within the resident microbiota and how its composition might impact their efficiency. Therefore, using the Botrytis cinerea – Pichia anomala strain K (well studied BCA) - apple pathosystem, our study aimed to answer two key questions: (i) what is the impact of the resident epiphytic microbiota on postharvest disease development and the efficacy of a BCA, and (ii) how can this impact be leveraged for limiting postharvest losses due to pathogens?

## Material and methods

### Sampling and plant materials

Apple fruits were harvested in September 2016 in Belgium from four different orchards (two commercial, one research, and one conservation orchards) just before harvest. Two *Malus sylvestris* accessions and 13 different cultivars of *M. domestica*, representing 18 samples in total, were sampled. More details are available in supplementary material 1.

### Microbiota harvest and biobanking

A pool of four fruit was considered as a sample replicate and 5 independent replicates were extracted per microbiota. The harvesting, biobanking, quantification, and preparation of microbiota is described in Supplementary Material 2.

### *In vivo* evaluation of disease control

The ability of microbial communities, isolated strains, and potential prebiotic molecules to control disease development by *Botrytis cinerea* Pers. was evaluated by standardised *in vivo* tests. The microbial communities (normalized at 10^5^ CFU/ml) were applied alone (n=10) or in combination with 10^5^ CFU/ml of *P. anomala* strain K (n=18) on sterilized commercial ‘Golden delicious’ wounded apples following standardized protocol (18) described in Supplementary material 3.

Eight prebiotic candidates were also tested *in vivo* following the same protocol at 0.5% and 1% (w:v) concentration with three selected microbiota. Twelve strains isolated from microbiota limiting the development of *B. cinerea* were tested *in vivo* at 10^7^ CFU/ml alone to evaluate their biocontrol efficacy.

### Taxonomical characterisation of the microbiota

Two biological replicates of each sample (18 microbiota x 2 replicates) were prepared in the laboratory and sequenced on Illumina MiSeq (GIGA, ULiège) according to the protocol described in Supplementary Material 4a.

For the bioinformatics analyses, demultiplexed paired-end FASTQ reads with trimmed adapters and primers were analysed using QIIME 2 (q2) version 2019-4 (19) and the SILVA_132 and UNITE version 8 taxonomic reference databases for bacteria and fungi, respectively. The detailed protocol is available in Supplementary Material 4a.

### Metabolic profile of the microbiota

The carbon metabolization profile of 17 microbiota (excepting Gb), of the BCA *P. anomala* strain K and of the pathogen *B. cinerea* strain V was assessed with commercial Biolog Ecoplates PM1 and PM2A microplate systems (Biolog, Hayward, CA, USA), containing 190 different carbon sources (20, 21) and following the protocol detailed in supplementary material 4. Each plate contained a negative control corresponding to sterilized water. The repeatability of the method was first checked using two aliquots of the same biological replicate of one selected microbiota called GoGx on a PM2A microplate (Figure Sup. Mat 4).

### Statistical data analysis

Comparisons of efficacy in controlling disease development during bioassays were made by the Analysis of Variance (ANOVA) at 5% type I error rate (α = 0.05) using RStudio version 3.5.1. Means of the normalized apple wound diameters were separated with the Tukey’s HSD test with the Agricolae 1.3-0 (22) R package. Corrected carbon source O.D data were scaled, co-clustered and Spearman correlations visualized in heatmaps with dendrograms using the R package Heatmap (23).

The impact of bacterial or fungal alpha diversity indices on the biological efficacy was estimated with Pearson correlation (α=0.05). In addition, Spearman rank pairwise correlations between all the biological efficacies (alone and at the three tested concentrations with the BCA), the taxa identified and the carbon sources metabolised were computed with the Microbiome R package (24) and plotted on R as heatmaps to drive the identification of beneficial strains and prebiotics of biocontrol. Because Spearman correlation investigates monotonic association rank and does not require parametric distribution, no distribution hypothesis was further assumed. Hence no p-values were inferred for later exploration.

### HTS-guided selection of beneficial strain and metabolism-based selection of prebiotics

Carbon sources or taxa whose metabolization or abundance showed a strong correlation with the biocontrol properties of the microbiota were evaluated *in vivo* to confirm their ability to support disease control as prebiotic or beneficial strains respectively.

The microbiota containing the highest abundance of targeted taxa were plated on PDA and R2A media, then each pure colony was purified twice and kept on the media used for isolation. Suspensions were prepared with sterile deionized water for taxonomy determination by PCR (Supplementary material 4b).

## Results

### Diverse microbiotas were harvested

The samples’ fungal and bacterial taxonomic composition were analyzed from 3,628,894 and 3,605,618 paired-end reads respectively. The reads were clustered into 1,916 bacterial and 5,442 fungal ASV, respectively. Analysis of beta diversity plots by Bray Curtis distance metrics for both bacterial and fungal ASV (Supplementary Material 5) showed that samples were well scattered but tended to cluster by location (PERMANOVA pairwise comparison pseudo-F test q-value < 0.05) [which included several factors like surrounding environment (plants, soil), management practices, and cultivars]. Among the 18 sampled microbiota, Proteobacteria (57 – 94%) and Bacteroidetes (2 – 24 %) were the most abundant bacterial phyla. The two major fungal phyla were Ascomycota (27 – 71%) and Basidiomycota (9 – 63%). Ten and 19 identified genera were present in more than 80% of the samples for bacteria and fungi, respectively, including seven of the eight genera considered as the core microbiota of apple surface. The observed percentages showed a high variability of taxa abundance (Supplementary material 6.a & b) between samples, but similar abundance between replicates. In addition, the average colony firming units (cfu) concentration of all microbiota samples was 2 10^4^ +/-1.49 10^4^ CFU / cm² of apple surface with a minimum of 2 10^3^ CFU/cm² (GoGx microbiota) and a maximum of 6.75 10^4^ CFU/cm² (M. *sylvestris* O57 microbiota). These concentrations were used as a proxy to normalise biocontrol assay (Supplementary material 2).

### The microbiota significantly impacted disease development

The ten microbiota tested alone provided a significant control (df=9, F=7.549, p=2.12 10e-10) of the disease caused by *B. cinerea* (strain V) with significantly different efficacies, ranging between 27% and 82% (Figure 1). One microbiota (P) provided a disease control significantly higher than the biocontrol strain K applied at the same concentration (10^5^CFU/ml) while seven microbiota (Rf, Po, Api, GoGx, GoTi, R48R0, Cer) presented an efficacy similar to the strain K. Two microbiota (JoTi and Ms) developed a lower protection than strain K.

**Figure 1.**
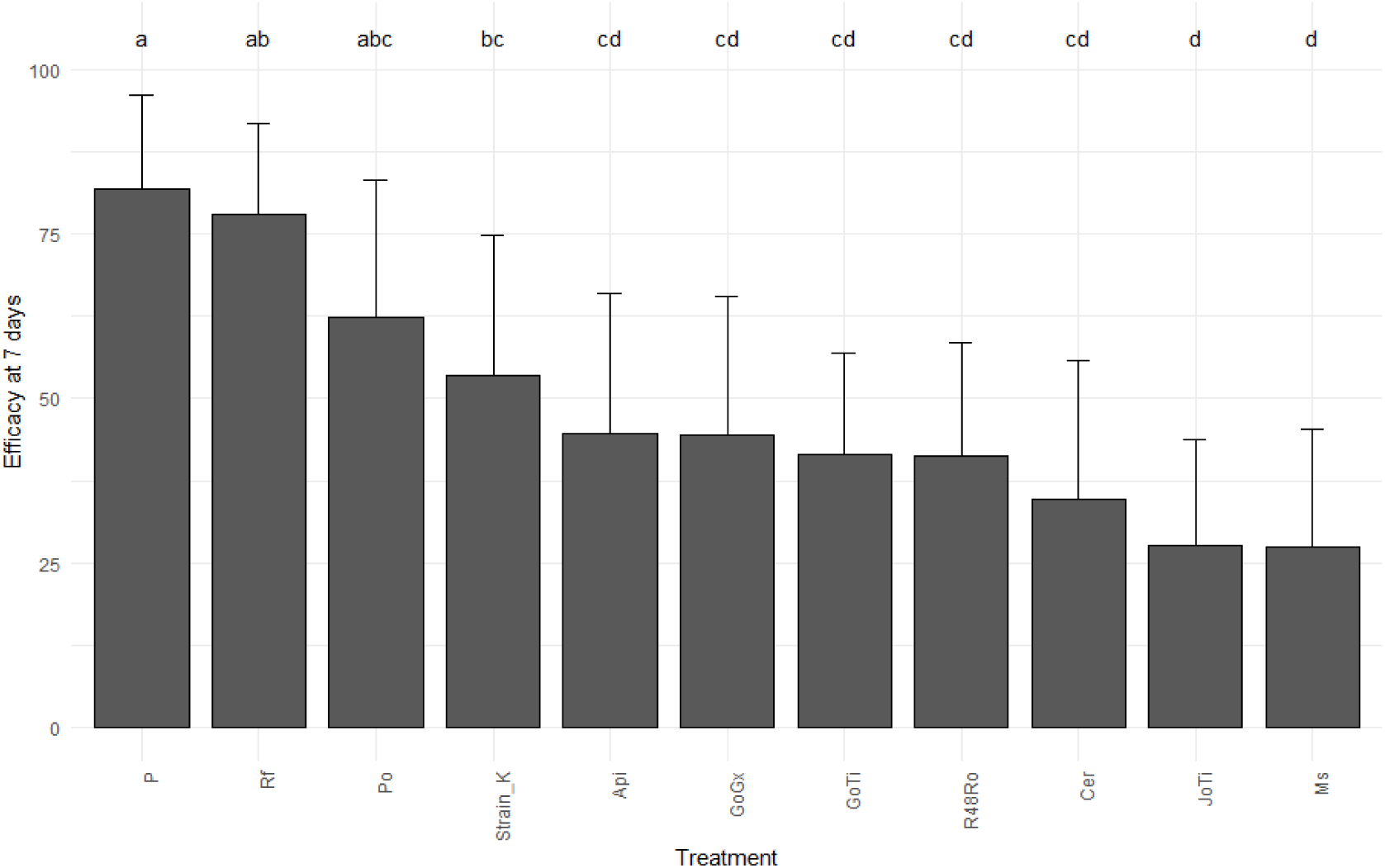
Efficacy of the apple microbiota (applied at 10^5^ CFU/mL; n=10) against *B. cinerea* rot development on apple wounds. Isotonic water was used as a control. Letters above each bar represent Tukey statistical mean groups. Error bars on each bar represent standard deviations.

### The microbiota modulated the efficacy of the biocontrol agent strain K

Overall, a highly significant impact of the microbiota on the strain K efficacy was observed (df=17, F=14.81, p value = 2 10e-16). The efficacy was improved by 100% for three microbiota (JoGx, O57 and Plp) or decreased by 27% with Ms (Figure 2). Importantly, the microbiota Cer, JoTi and R48Ro whose efficacies alone were very low, significantly raised the protection against *B. cinerea* when applied with strain K.

**Figure 2.**
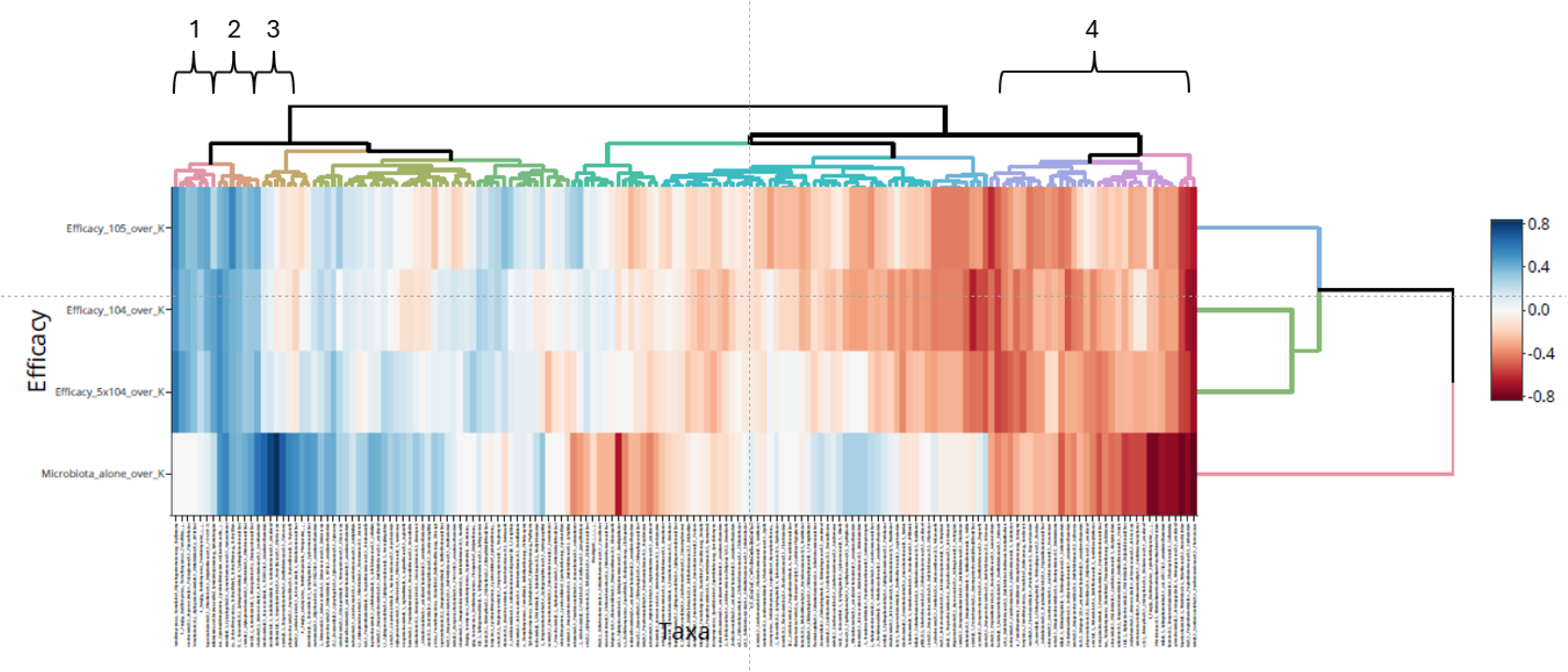
Clustered heatmap of correlations between biological efficacy and taxa abundance in the studied microbiota. Each color represents a correlation, and the strength of the correlation is showed at right side in the legend. Each column represents microbial assigned taxa and each line represents the efficacy of biocontrol assay at each concentration. Blue and red colours indicate positive and negative correlations, respectively.

The impact of the microbiota was also tested at two other concentrations: 10^4^ and 5x10^4^ cfu/ml (Supplementary Material 7). An interaction effect between the concentration and the microbiota samples was observed (F=2.369, df=34, p< 0.001) but concerned only some microbiota. For example, the 3 microbiotas presenting the highest efficacy increase (+100%) over 10^5^ cfu/ml maintained their level of efficacy at the two other concentrations.

### The microbiotas are contrasted carbon metaboliser

The repeatability of the test was suitable as the prior evaluation of two biological replicates of GoGx did not show any significant difference in their metabolization profile with the lowest difference observed after 24h (Supplementary material 4). The seventeen microbiotas (all but Gb microbiota whose stock was empty) showed contrasted profiles of carbon source metabolization and were able to metabolize 186 carbon sources with high variability profile (Supplementary material 8). Among all the carbon sources tested, 19 were metabolised by at least 12 microbiota (70 %). No carbon source was metabolised by all the microbiota tested, but the most metabolised were D-galactose and laminarin. The 17 microbiota and the 3 isolated strains were clustered according to their metabolization pattern (Supplementary Material 8). The largest cluster included 15 microbiota. Two microbiota (Plp and M) were distant from the others.

### The abundance of some taxa is correlated with disease protection, whatever the microbiota

First, bacterial or fungal alpha diversity indexes were not correlated with the efficacy of apple microbiota applied alone or with the BCA. On the other hand, Spearman correlations between biological efficacy (alone and at all concentrations) and taxa abundance identified clusters positively or negatively correlated with the biocontrol activity with or without the strain K. (Figure 3). Cluster 1 corresponded to taxa abundant in microbiota such as members of *Drechslera, Galactomyces, Chitinimonas* or *Acaulospora* that were associated with biocontrol when co-applied with strain K. Cluster 2 included taxa such as *Kondoa*, *Neofabrae* and *Hymenobacter*, whose abundance in microbiota was correlated with a high protection for the microbiota applied with strain K or alone. Cluster 3 corresponded to taxa, such as *Rhodococcus*, *Variovorax*, and *Polaromonas* that were correlated with high microbiota protection when applied alone but not when the strain K was co-applied, suggesting antagonism. clusters were also of interest. The cluster 4 included taxa, belonging to *Ralstonia*, *Janibacter*, *Erwinia*, *Cutibacterium* and *Aeromicrobium* genera, that were more abundant when the microbiota strengthened the disease severity applied alone or with the strain K. Detailed table of these taxa with their correlation values are available in Supplementary Material 9a.

**Figure 3.**
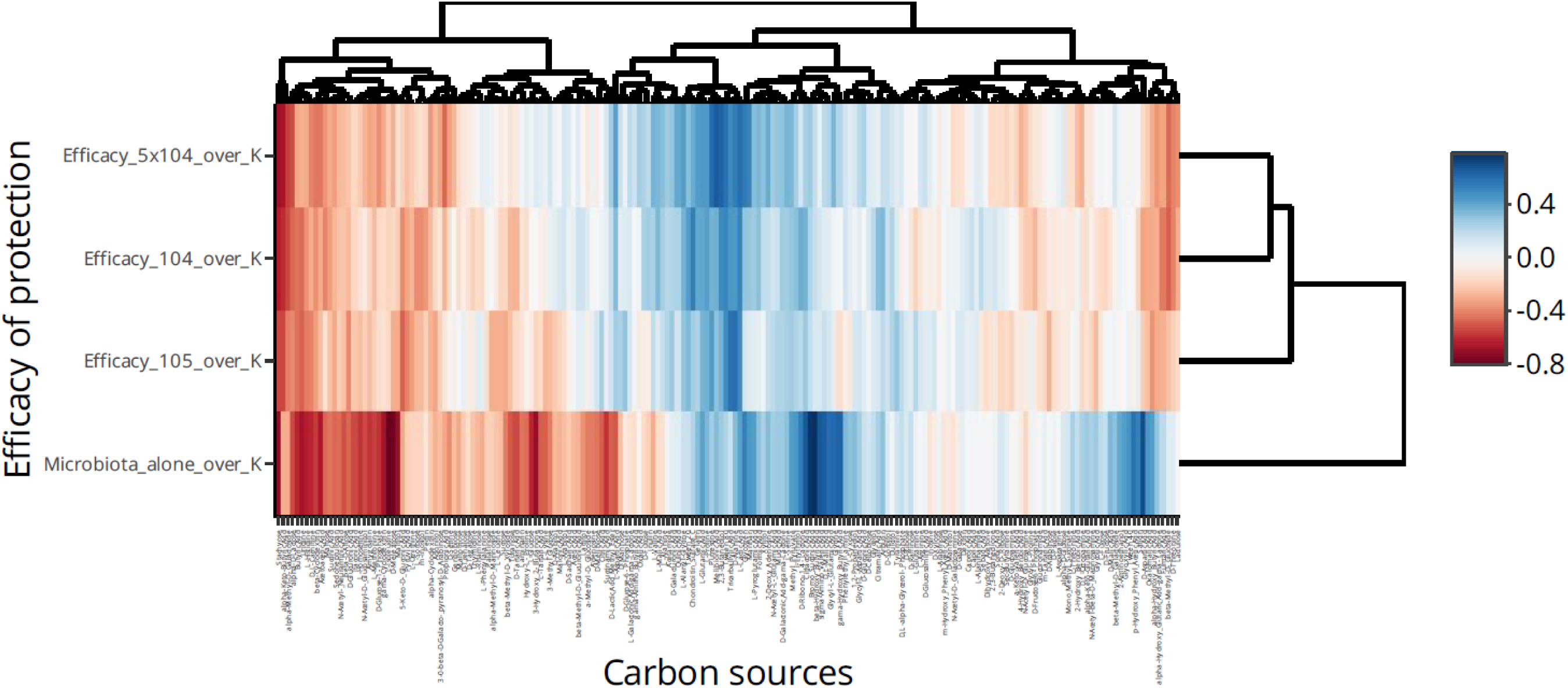
Clustered heatmaps of correlations between biological efficacy of microbiotas (applied alone or co-applied at 3 concentrations with strain K) and the efficacy of carbon source metabolization for the studied microbiota. Each column represents a carbon source and each line represents the efficacy of biocontrol assay at each concentration. Each colour represents a correlation, and the strength of the correlation is showed at right side in the legend. Blue and red colours indicate positive and negative correlations, respectively.

### The metabolization of some carbon sources is correlated with disease protection

For the carbon source, the metabolization of Glycyl-L-Proline, D-Fructose-6-_Phosphate, D-Arabinose, L-Asparagine, alpha-D-Glucose, D,L Malic acid, L-Threonine, Glycolic_Acid, L-Fucose and D-Fructose was positively correlated with the efficacy of the microbiota applied alone (Figure 4). In addition, the metabolization of D-Saccharic_Acid, Acetic_Acid, beta-Methyl-D-_Glucuronic_Acid, Lactitol, gama-Amino_Butyric_Acid, L-valine and Inulin was correlated with the efficacy of strain K. Meanwhile, higher metabolisation of molecules such as Xylitol, 2-Deoxy_Adenosine, N-Acetyl-L-_Glutamic_Acid, L-Serine, Methyl_Pyruvate, i-Erythritol, L-Proline and D-Alanine was positively correlated with the protection of strain K or the microbiota. For another cluster of molecules (m-Inositol, Pectin, L-Isoleucine, Maltitol, Maltose, Butyric_Acid and L-Methionine), their higher metabolization was correlated with a lower control of *B. cinerea* by the strain K or the microbiota. Detailed table of these carbon sources correlations values are available in Supplementary Material 9b.

### Selection and *in vivo* evaluation of prebiotics for disease control

Eight molecules were selected based on the correlation of their metabolization by the microbiota and the disease control brought by the microbiotas but also on their availability, cost and presence in edible food. Their prebiotic potential was tested with strain K and with a microbiota showing middle to low protection (GoGx). The protection achieved with GoGx microbiota applied alone or with strain K were similar to the obtained efficacies during the initial evaluation of GoGx efficiency. The impact of prebiotic molecules applied with GoGx on *B. cinerea* pathogenesis was variable with efficacy ranging from 91% to 4%. L-alanine and Meso-erythritol were the most efficient to improve the disease control of GoGx + strain K with 3.6X and 4.9X wound diameter reduction, although with high variability (Table 1). These two molecules and the xylitol were also tested *in vivo* (at 0.5%) with GoGx or with one of two other microbiotas (Api and GoTi, belonging to the same statistical group as GoGx in term of efficacy) and with strain K (figure 4). The strain K alone achieved a protection of 51 % (C.V. = 41%) compared to the control. Whatever the microbiota, the addition of L-alanine did not raise the disease protection compared to strain K applied alone, including when applied with GoGx, indicating a variable effect throughout time. The application of meso-erythritol or xylitol to Api and GoGx microbiota significantly improved the protection compared to strain K alone but no difference was observed when added to GoTi.

**Table 1:** Evaluation of the prebiotic properties of 8 molecules when applied (at 0.5%) with GoGx microbiota (and strain K on the biocontrol of *B. cinerea* through an *in vivo* bioassay on apple.

| Treatment | Molecules | Lesion diameter (mm) | Standard deviation (mm) | % of protection | Tukey's statistical analyses |
| --- | --- | --- | --- | --- | --- |
| Microbiota + strain K + | L-alanine | 1,6 | 2,5 | 91% | d |
| Microbiota + strain K + | Meso-erythritol | 2,2 | 2,9 | 87% | d |
| Microbiota + strain K + | L-Threonine | 6,7 | 4,7 | 60% | cd |
| Microbiota + strain K + | - | 7,9 | 4,9 | 54% | bcd |
| Microbiota + strain K + | Xylitol | 8,0 | 5,4 | 53% | bcd |
| Microbiota + strain K + | GABA | 8,8 | 7,7 | 48% | bc |
| Microbiota + strain K + | 2-deoxy-D-Ribose | 10,8 | 7,6 | 36% | abc |
| Strain K | - | 11,2 | 3,2 | 34% | abc |
| Microbiota | - | 11,9 | 6,1 | 30% | abc |
| Microbiota + strain K + | L-proline | 14,3 | 12,6 | 15% | ab |
| Microbiota + strain K + | Gluconic acid | 16,3 | 10,7 | 4% | a |
| Isotonic water | - | 16,9 | 5,6 | n.a. | a |

### Selection of beneficial fungi and *in vivo* evaluation of their efficacy

The three best microbiota (JoGx, O57 and Plp) were selected for their biocontrol properties and their potential higher relative abundance in fungal taxa identified as beneficial. After plating and isolation, 125 strains were isolated, purified, and identified by sequencing their ITS. Many isolates belonged to known BCA genera (*Aureobasidium, Debaryomyces, Metschnikowia, Rhodotorula,* and *Sporobolomyces*) but none of them belonged to the genera highlighted in the beneficial clusters 1, 2 and 3. On the other hand, two isolated strains belonged to species (*Farysia acheniorum* and *Kalmanozyma fusiformata*) included in genera never mentioned in biocontrol. Evaluation of beneficial properties was carried out by *in vivo* biocontrol assay on 12 isolates. *In vivo* assays showed that all the isolated strains provide a significant protection against *B. cinerea* rot (df=11, F=3.541, p= 9.89 10e-05) with an efficacy ranging from 93 to 61 % (Supplementary material 10). Two strains of *Metschnikowia pulcherrima* and the strain of *Farysia acheniorum* controlled disease development by at least 90%.

## Discussion

### The impact of resident microbiota on disease development is variable and could rely on a complex network of interactions

Microbial communities of apple fruit can present attractive, functional biocontrol traits and genes (25), which can be harnessed for the ecological management of postharvest pathogens. Except hypotheses casted in the literature, there has been no evaluation of the impact of epiphytic microbiota on fruit disease development. In this publication, we demonstrated the contrasted impact of a diverse set of microbiotas from apple carposphere on disease development and its biocontrol.

First, the contrasted taxonomic profiling of the harvested microbiota showed that the most abundant fungal and bacterial phyla and genera were similar to previous studies on apple fruit (11, 26), and included genera of pathogenic species or of biocontrol agents.

The contrasted microbiotas exhibited variable levels of protection against *B. cinerea*, ranging from 30% to 100%. In our bioassay, the protection, obtained under standardized *in vivo* conditions, primarily depended on the complex set of trophic interactions between the pathogen and the members of the microbiota. Noteworthy, the pathogenesis of *B. cinerea* was never reinforced. This phenomenon could be attributed to competition for available nutrients and a niche occupation by microbiota members, limiting the pathogen development.

In addition, the microbiotas and, in some cases, their concentration impacted the biocontrol efficacy of the strain K which either raised (up to double) or dropped (down to 27%). This contrasting impact recalled the variability of BCA efficacy in real conditions and suggests that the BCA efficacy is strongly influenced by the variability of ecological relationships occurring with the resident microbiota in the processes of biocontrol phenotype expression. Interestingly, three microbiotas raised the efficacy of the BCA strain K while providing only a low protection when applied alone. They may have a synergistic interaction with strain K and could be a valuable source of helper strains (27).

The demonstration of the impact of resident microbiota on disease development and biocontrol strain efficacy opens an innovative research area to understand this phenomenon. Interaction models (28) and omics technologies (29) used to decipher in situ BCA-pathogen interaction could be adapted to the study of microbial communities within a proper ecological framework (1).

### Which component of microbiota alpha diversity is important for pathogen control?

Our results represent the first interventional study on fruits where resident microbiota are harvested and further tested for biocontrol properties. A higher richness or evenness was not associated with the higher biocontrol efficacy of apple fruit microbiota against *B. cinerea* while most observational studies comparing healthy and diseased plant material suggested that high alpha bacterial diversity promoted disease control (30) (31) (32) (33). A first explanation of this apparent contradiction is that our results are unique as they are based on standardized bioassays with microbiota application, not on observations prior or after disease development. Our results uncover the complexity of interactions: beyond the diversity of the microbiota, the crucial factor could be the active members of the network developing biocontrol properties and their ecological interactions. This complexity was also recently suggested by a large-scale observational study on the association between *Arabidopsis* leaf- and root-associated microbiota and disease development (34).

### Microbiome-guided selection of beneficial microorganisms by HTS taxonomic profiling can be strengthened by functional characterization of microbiota but remains a challenge

An HTS-guided strategy can be applied to select beneficial microorganisms whose presence or abundance in microbial communities is associated with beneficial properties. This concept is recent and has only been applied on lettuce(5), grapevine(35), citrus(36) or *Coptis chinensis*(37), *A. mongholicus* (4) and *A. thaliana* (34). Theoretically, this selection drastically reduced the number of isolated strains to be tested compared to empirical methods. Nevertheless, this strategy has biological and technical limitations. First, microorganisms present at very low proportions can play a fundamental role in the microbiota functions (38), but their low proportion, often accompanied by higher variability, will limit their probability of being highlighted and selected. In addition, if the comparison is carried out between microbiota of healthy and diseased tissue, it can introduce a bias as the microbiota modification on diseased tissue might be a consequence of disease development without influencing the first stages of disease initiation. Our approach of microbiota application before pathogen inoculation limited this bias although the taxonomic composition of the microbiota might have changed between the application on the wounds and the application of the pathogen 24 hours later Another challenge is linked to biology of the species, strain isolation protocols targeting specific taxa might be missing as some taxa could be difficult to isolate due to the lack of information on their ecology or impossible to culture under the current knowledge. One way to tackle this challenge might rely on culturomics technics (39).

During this study, we could not isolate strains of the genera identified in beneficial clusters, confirming the difficulty of HTS-guided selection. This challenge was compensated by another approach relying on the isolation of strains from the most beneficial microbiota. Most isolated taxa were already mentioned in the literature for their biocontrol properties but, a strain from *F. acheniorum,* a species never cited as biocontrol agent on plants biocontrol properties. It should be better characterized for its biocontrol properties and metabolism to evaluate its compatibility with existing biocontrol strains, for example, within synthetic communities. Our approach was *in fine* guided by biocontrol properties of the microbiota and showed interest for isolating a more diverse set of putative beneficial microorganisms.

### Engineering the microbiota for disease control by targeted selection of prebiotics molecules

The functional catabolic diversity of microbiota can be evaluated by studying their ability to metabolize various sources of nutrients. Such functional characterization can lead to the identification of prebiotics of biocontrol (15). These molecules are substrates that can modulate the microbial composition of the microbiota toward an improved control of disease development. For instance, tagatose has been tested as a prebiotic of biocontrol against grapevine leaf downy and powdery mildew, causing a shift in the leaf microbiota (40). Glutamic acid has also reshaped tomato rhizosphere by increasing the abundance of several taxa while, at the same time, reducing disease symptoms (41). Correlating the metabolic profile of microbiota with their biocontrol properties when applied alone or in the presence of strain K identified several clusters of potentially valuable molecules with high Spearman correlations with the biocontrol properties. Eight of these molecules were further tested in bioassays including a microbiota, the strain K and/or the pathogen. The results were variable (from -50% to +37% in efficacy) depending on the molecule. This observation suggested that the large-scale correlations carried out should be considered as a preselection and underlined the need to confirm *in vivo* the candidate molecules. The L-xylitol and Meso-erythritol increased the efficacy of the microbiota alone, and in the presence of strain K. These two molecules are rare sugars at low concentrations in apple fruit. Rare sugars have been shown to impact pathogen development and biocontrol of BCA. For instance, the rare sugar D-tagatose inhibited the growth of pathogens like *Phytophthora spp.* (42) and increased the efficacy of BCA on many fruits and vegetables (43). L-xylitol and meso-erythritol correspond to the most promising biocontrol molecules in this pathosystem and deserve further characterization of their mode of action. Indeed, meso-erythritol was not well metabolized by any of the 3 tested microbiota (Api, GoGx, GoTi) and the two pathogenic strains of *Botrytis*, but was the highest molecule metabolized by the strain K. This can explain why this molecule had increased the biocontrol activity in the presence of the strain K. Without the strain K addition, this molecule may have promoted other beneficial microorganisms toward a biocontrol phenotype. On the other hand, another selected carbon source, L-alanine, was reported to increase the efficacy of BCAs on apple (44, 45) and pear (46).

Overall, as suggested by our results, the targeted selection of molecule could be envisioned as part of the biocontrol strategy against plant pathogens, applied alone to modify the resident microbiota or applied with BCA applied alone or in synthetic communities to foster the expression of their biocontrol phenotype. Beyond our exploratory study, a broader screening of molecules, preferably recognized as GRAS, could be envisaged to improve the field efficacy of biocontrol agents applied alone or in synthetic communities.

In conclusion, this study shows that apple fruit epiphytic microbiotas significantly influence postharvest disease development. The microbiotas’ taxonomic and metabolic profiles correlated with biocontrol evaluation identified potential biocontrol prebiotics and beneficial strains. These results highlight the importance of the pathobiome concept and underline that residing microbiota is an essential component to be considered when developing solutions to control postharvest diseases.

## Notes

### Competing Interest Statement

The authors have declared no competing interest.

### Summary of Updates

The text has been optimized/rationalized and the clustering figures are now provided in high definition, allowing their analysis at high resolution

