## Supplementary Material for "Highlighting the complexity of pathogenesis: the host microbiota impacts disease development on apple fruit and is a cornerstone for its biocontrol"

**Supplementary material 1: Beta-diversity analyses for the harvested microbiotas**

**Figure Sup. Mat. 1:** Beta diversity plots of Bray Curtis distance metrics coloured by sampling location. Each location represents diverse cultivars under contrasted management practices: Organic = commercial orchard, organic farming, CONV = commercial orchard, integrated pest management, NT = germplasm conservation orchard; Light_org = research orchard with reduced spraying

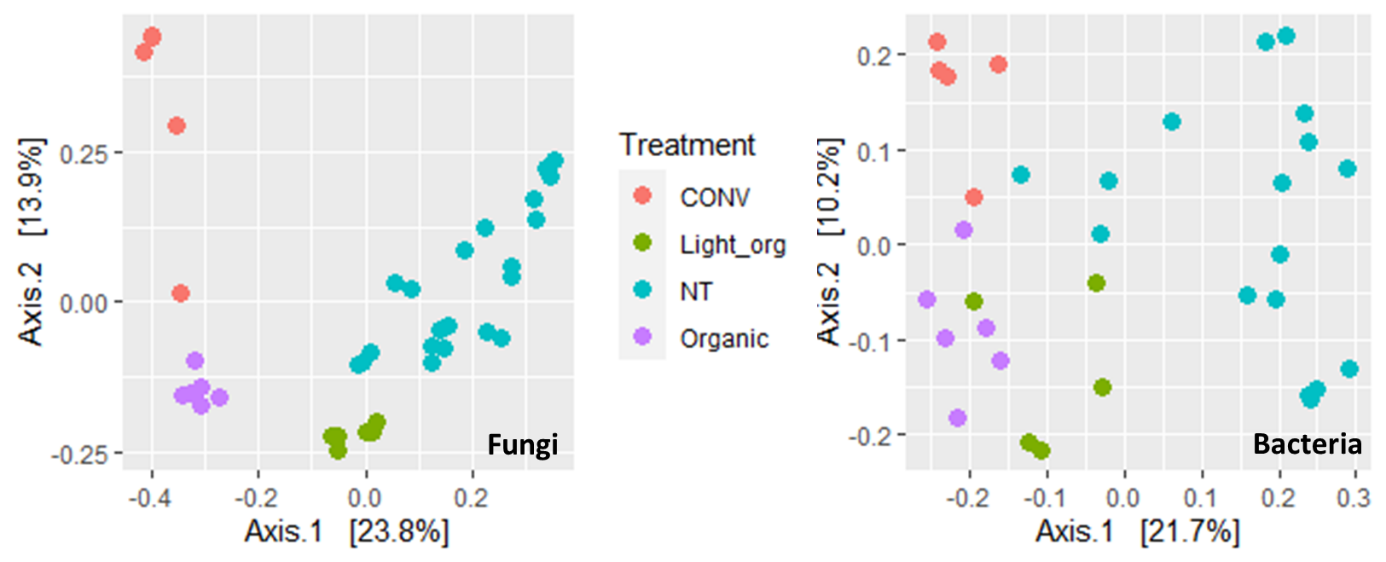

**Supplementary material 2: Taxonomy composition and alpha diversity indexes of harvested microbiotas (Raw data available on SRA: Accessions PRJNA663271 to PRJNA662931)**

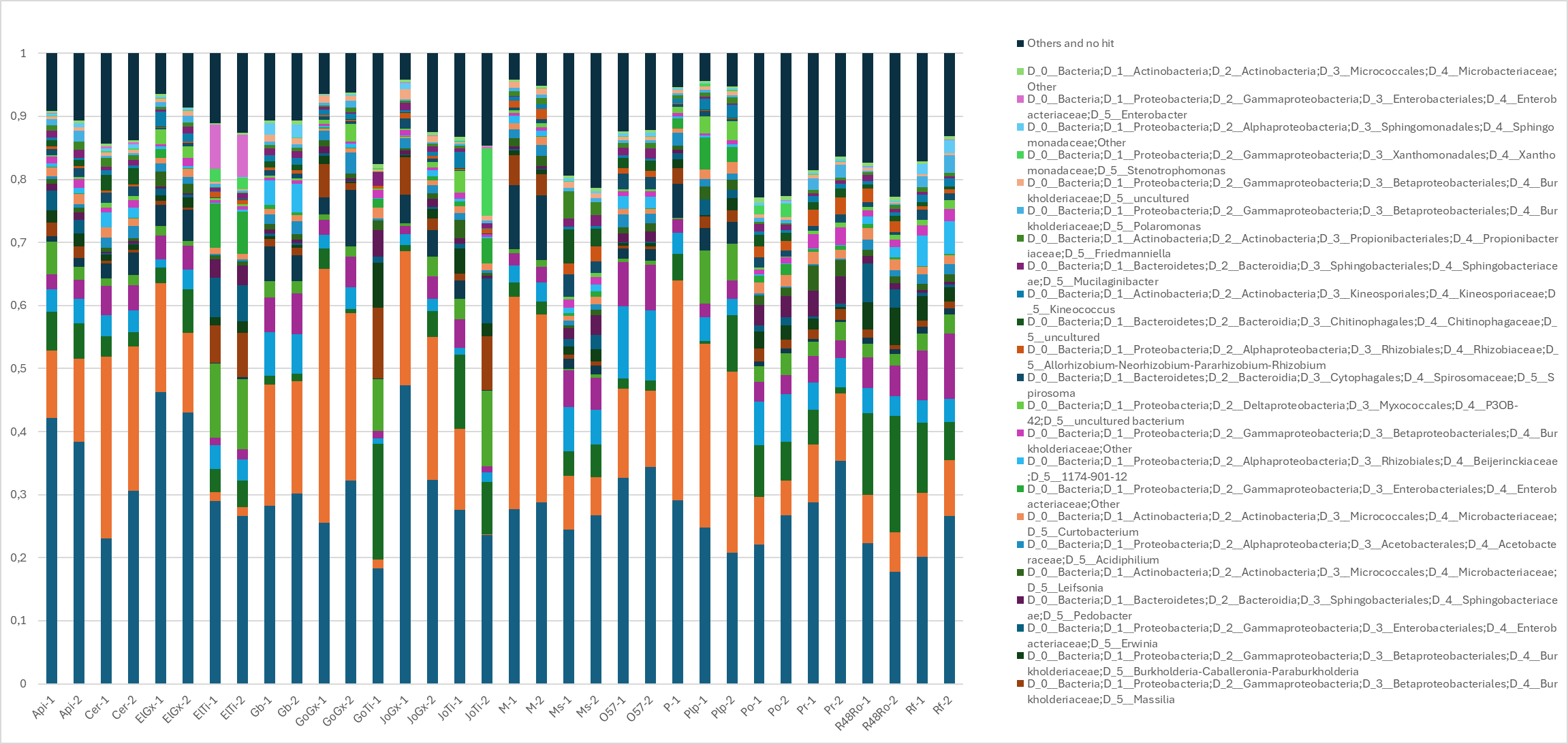
**Figure Sup. Mat 2a.** Overview of the bacterial taxonomic profile at genera level for the microbiota harvested from apple fruit carposphere (two replicates per sample, except for GoTi and P, read normalisation depth of 29,666). Each bar corresponds to a sample and each colour represents a taxa whose size is proportional to its relative frequency. Only the 30 most abundant bacterial tax are displayed.

**Figure Sup. Mat 2b.** Overview of the fungal taxonomic profile at genera level for the microbiota harvested from apple fruit carposphere (two replicates per sample, except for JoTi and P, read normalisation depth of 1,028). Each bar corresponds to a sample and each colour represents a taxa whose size is proportional to its relative frequency. Only the 20 most abundant taxa and two replicates per microbiota are displayed

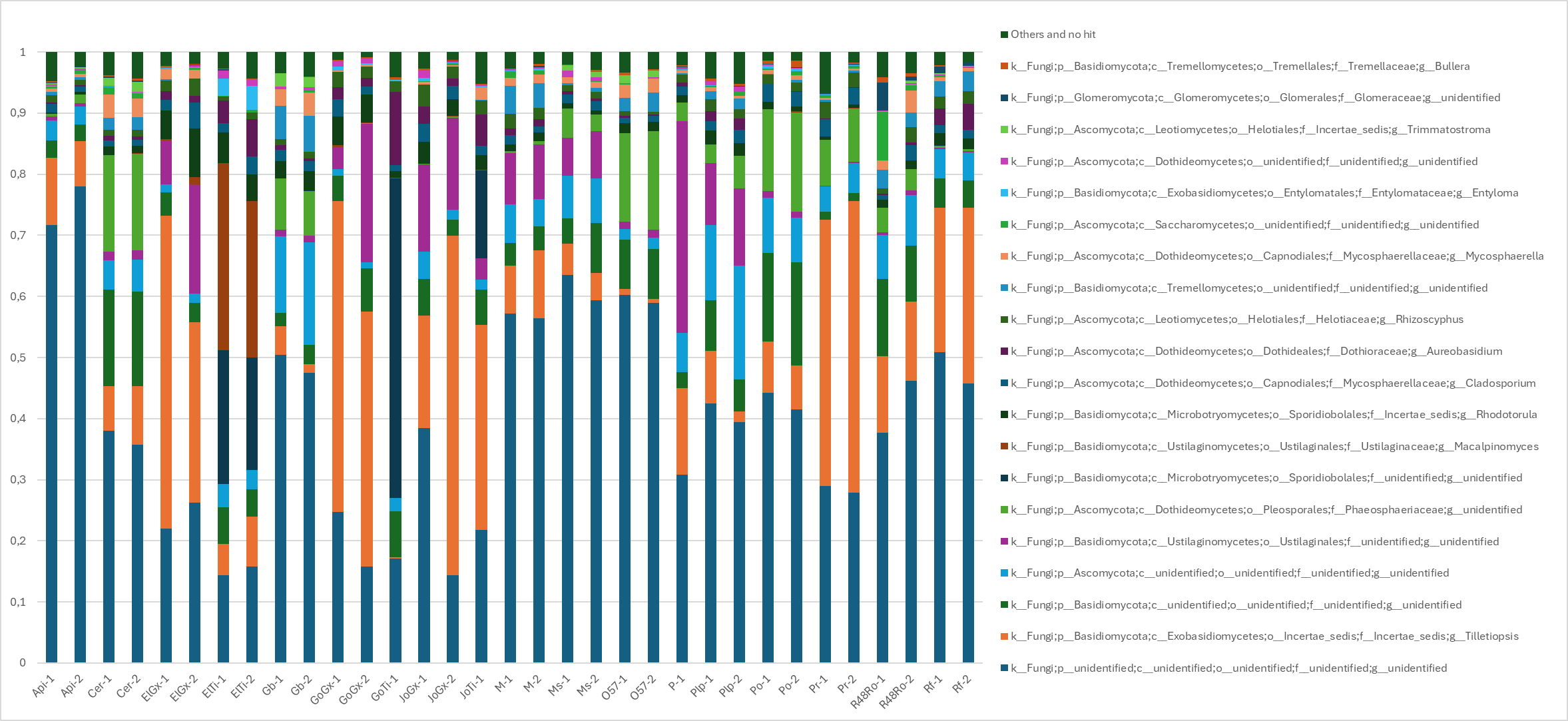

**Supplementary material 3. Detailed protocol on biocontrol in vivo bioassay**

Three wounds of 6 mm diameter and 2mm depth were made per apple with a cork-borer. Forty microliter of inoculum suspension was applied per wound and 10 fruits were used per treatment. A negative control (8.5% NaCl water) was used in all the experiments, as well as the strain K alone at 10^5^ CFU/mL as reference. Two other concentrations of microbiota (10^4^ CFU/ml and 5 x 10^4^ CFU/ml) were also tested in co-application with the strain K to build a robust database for correlation analysis. After 24h, wounds were inoculated with 10e6 spores/ml of *Botrytis cinerea* strain V or *Penicillium expansum* strain 880. The rot diameters were measured after 10 days and the 6 mm diameter of the cork-borer was subtracted from each measure. The diameter of the diseased lesion recorded on each wound was further normalized on the strain K efficacy or the negative control depending on the comparison made with the formula:

$$Efficacy \%=\frac{Diameter of control (or strain K) - Diameter of treatment}{Diameter of control (or strain K)}$$

**Supplementary material 4**. Impact (in percentages of efficacy modification) of the application of apple microbiota (n=18) on the biocontrol efficacy of strain K against *B. cinerea* V on apple wounds. Letters above each bar represent Tukey statistical mean groups. Error bars on each bar represent standard deviations. The strain K applied alone reached an efficacy of 52% and corresponded to 0% in this graph.

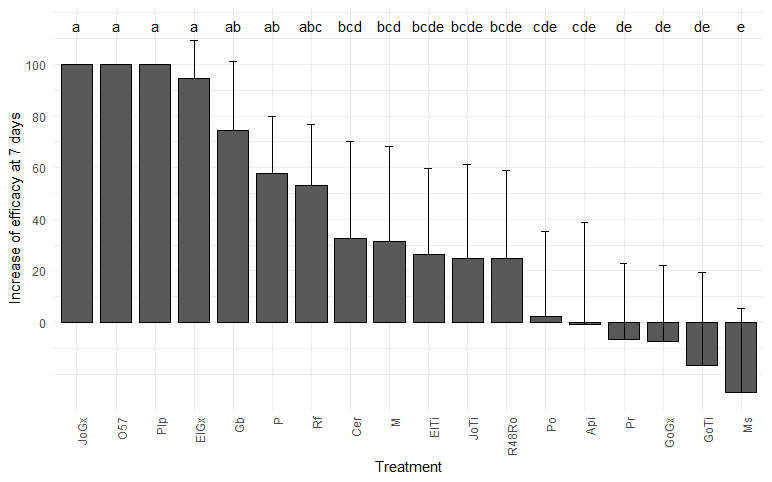

**Supplementary Material 5: Impact (in %) of 18 microbiotas applied at 3 concentrations on the biocontrol efficacy of strain K against *Botrytis cinerea***

**Figure Sup. Mat. 5:** Efficacy (in %) of 18 microbiotas applied at 3 concentrations on the biocontrol efficacy of strain K against *Botrytis cinerea*. All efficacy data were first normalized on the efficacy of the strain K applied alone at 10^5^ CFU/ml. A=10^5^ CFU/ml; B= 5 x10^4^ CFU/ml; C= 10^4^ CFU/ml.

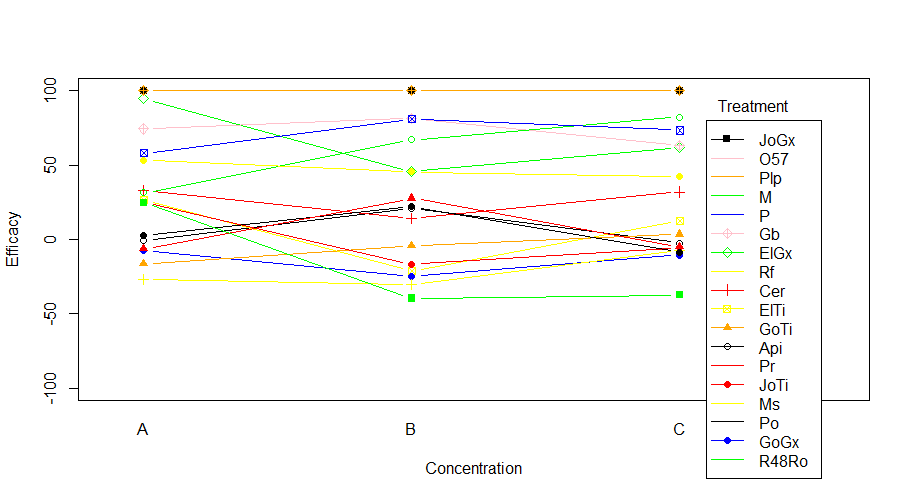

**Supplementary material 6 : Metabolic profiling of the microbiotas**

One hundred microlitre of microbial suspension, normalized at 10^5^ CFU/ml, in sterile 0.85% NaCl solution with 1% (120µl in 12 ml total microbial mix solution) of the tetrazolium redox dye H was loaded in each microplate well, and microplates were stored at 25°C in darkness. Optical Densities (O.D) at 492 mm and 595 nm wavelength were recorded twice at 10s interval (two technical replicates) with a 96 well plate Spectrophotometer Multiskan Ascent (Thermo Labsystems, Helsinki, Finland). Optical Densities (O.D) at 492 nm wavelength was recorded at T0 and each 24h up to 7 days. The results of the repeatability test suggested that 24 h was prefered for downstream analysis because at this moment, the smallest variability between replicates during the repeatability tests was recorded.

Technical replicates were similar (higher variation for all wavelength =0.01mm), but the downstream analysis was done with 492 nm (estimated measure of cell respiration) data after 24 h (very low difference between replicates).

For each time point and well, the mean of the two raw O.D values (XiA, XiB) were corrected before data analysis and the formula used was: Xi = ((Xi_24_ – Xi_0_) – Ci – S.D_24_), where: i= carbon source; Xi_0_ (Xi_24_) = O.D of i at 0h (after 24 h) and is the average of XiA and XiB; Ci= O.D negative control at 24h – O.D negative control at 0h (Corrected O.D of the negative control of the plate containing i); and S.D_24_ = 3 x S.D. (standard deviation of all the negative control in all plates). O.D. values remaining negative after the correction were not set to zero for the microbiota samples because certain types of microorganisms are capable to secondly reduce the formazan obtained after the first reduction of the tetrazolium dye. The results are summarized in figure Sup. Mat 6.

**Figure Sup. Mat 6**: Average of optical densities obtained with two replicates (R1 and R2) of the GoGx microbiota for 190 carbon sources every 24 hours during 7 days. a) optical density at 492 nm; and b) optical densities at 595 nm

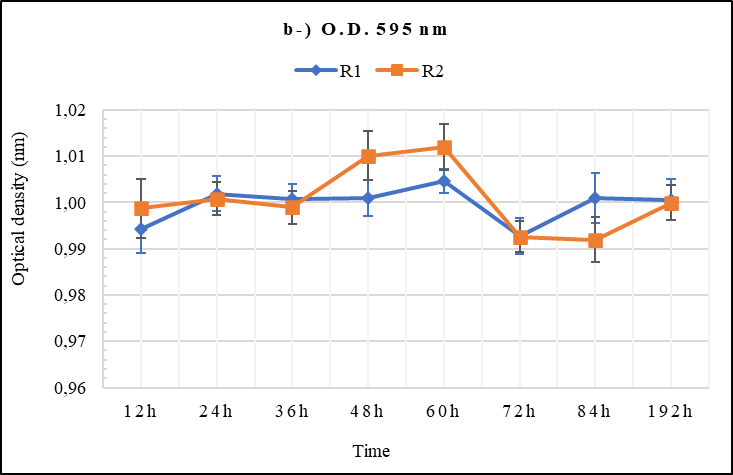

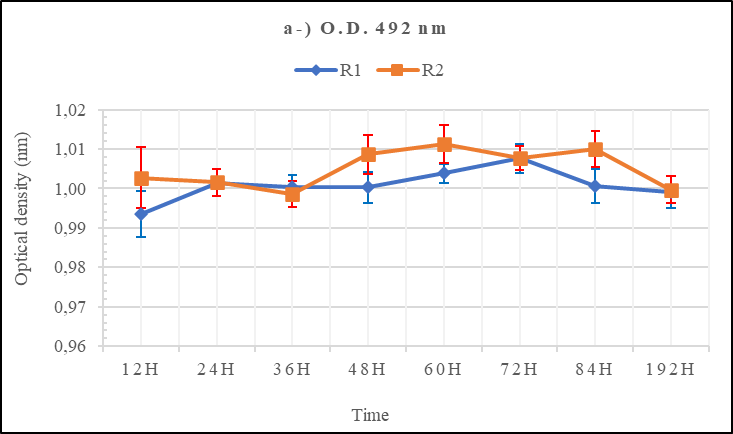

**Supplementary material 7.** **Co-clustering analysis of the metabolization of 190 carbon sources by 17 apple fruit microbiota**

**Figure Sup. Mat. 7**: Co-clustering analysis of the metabolization of 190 carbon sources by 17 apple fruit microbiota samples, *Pichia anomala* strain K, and two strains of *Botrytis cinerea* (DCS and V).

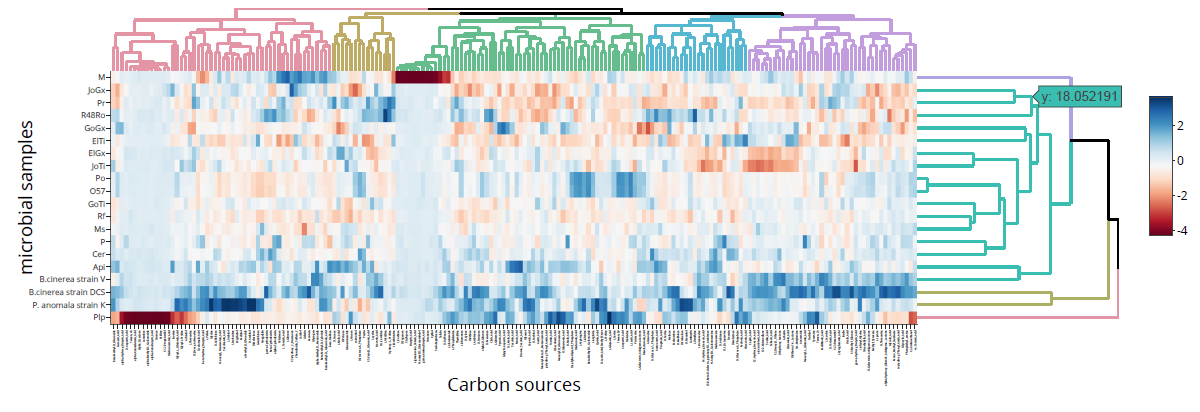

**Supplementary material 8a. (separate excel file)** Detailed table of Spearman correlation between taxa identified by metabarcoding and the efficacy of microbiotas applied alone or at 3 concentrations (10e4, 5x10e4, and 10e5) with the strain K (applied at 10e5)

**Supplementary Material 8b. (separate excel file)** Detailed table of Spearman correlation between carbon metabolisation intensity and the efficacy of microbiotas applied alone or at 3 concentrations (10e4, 5x10e4, and 10e5) with the strain K (applied at 10e5)

**Supplementary material 9.** Evaluation of the prebiotic potential of L-Alanine (Ala), meso-erythritol (Ery) and xylitol (Xyl) applied at 0.5% with 3 microbiota (Api, GoGx, GoTi) and the strain K against disease development by B*. cinerea*. The control in orange corresponds to the strain K applied alone. Letters for each bar represent Tukey statistical mean groups. Error bars on each bar represent standard deviations.

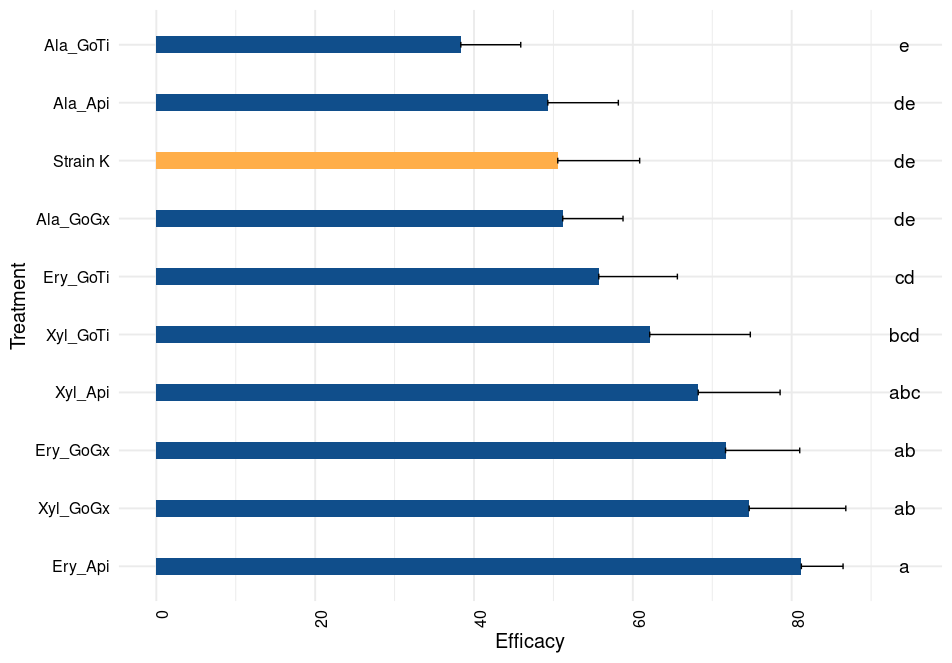

**Supplementary Material 10**: biocontrol activities of fungal strains isolated from microbiotas

**Figure Sup. Mat. 10.** In vivo biocontrol efficacy against *B. cinerea* of 12 isolated yeast (applied at 10e7 ufc per wound) expressed as lesion diameter (in cm) compared to non-treated control. Letters above each bar represent Tukey statistical mean groups. Error bars on each bar represent standard deviations.

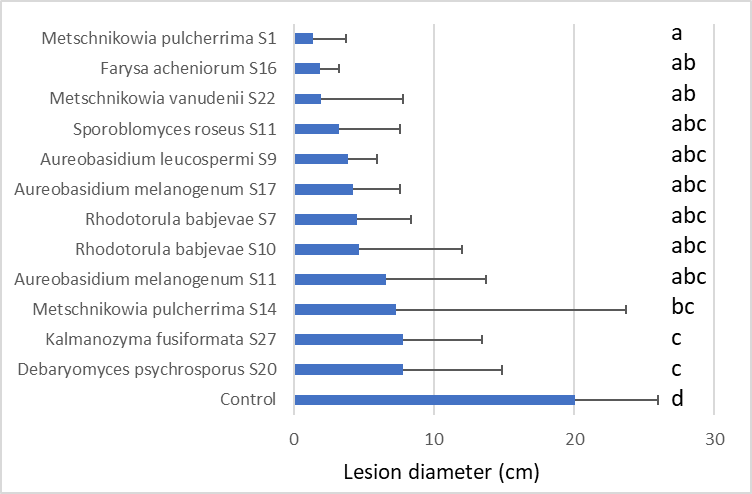

**Supplementary material 11.** **Details on the sampled apple fruit for epiphytic microbiota harvest**

Orchards corresponded to two commercial orchards managed under organic farming (Site A - 50°33'47.5"N; 4°43'17.8"E) or under Integrated Pest Management (IPM), (Site B - 50°48'36.7"N; 4°57'08.8"E), one 35-years old conservation orchard never treated with pesticides from the Fruit Tree Genetic Resources Collection of Walloon Agronomic Centre of Gembloux (Site C – CRA-W - 50°33'47.5"N; 4°43'17.8"E) and a research orchard of CRA-W combining organic farming with reduced spraying of copper (Site D - 50°33'06.8"N; 4°39'11.0"E).Trees from CRA-W were grafted on M9 rootstock with 4 m x 2 m in the conservation orchard (site C) and 4 m x 1.5 m in the light organic orchard (site D). M9 rootstocks were produced with a spacing of 3.5 m x 1.25 m in commercial orchards. Light organic orchard was treated with reduced doses of copper (0.5 kg / ha, 3 times), wettable sulfur (6 kg / ha, 6 times) and Sulfocalcic porridge (8 kg / ha, 3 times). Fruits were harvested each time on one three with new sterile gloves and collected in sterile plastic bags. In total, 18 different apple samples were collected (Table 1).

**Supplementary table 11.** Species, cultivar names and sampling site for the apple fruits sampled for biobanking and characterizing their epiphytic microbiota. Sampling sites: A = commercial orchard, organic farming, B = commercial orchard, integrated pest management, C= germplasm conservation orchard; D = research orchard with reduced spraying

| **Species** | **Cultivar** | **Sampling site** | **Code** |
| --- | --- | --- | --- |
| *Malus domestica* | Api Etoilé | C | Api |
| *M. domestica* | Ceres | C | Cer |
| *M. domestica* | Elstar | A | ElGx |
| *M. domestica* | Elstar | B | ElTi |
| *M. domestica* | Golden Delicious | A | GoGx |
| *M. domestica* | Golden Delicious | B | GoTi |
| *M. domestica* | Gris Braibant | C | Gb |
| *M. domestica* | Jonagold | A | JoGx |
| *M. domestica* | Jonagold | B | JoTi |
| *M. sylvestris* | N.A. | C | Ms |
| *M. domestica* | Marnica | D | M |
| *M. sylvestris* | N.A*.* | C | O57 |
| *M. domestica* | Patte de Loup | D | Plp |
| *M. domestica* | Pinova | D | P |
| *M. domestica* | Pomme de Dame | C | Po |
| *M. domestica* | Président Van Dievoet | C | Pr |
| *M. domestica* | La Rossenfosse | C | R48Ro |
| *M. domestica* | Reinette de Flandres | C | Rf |

**Supplementary Material 12. Additional details on microbiota harvesting protocols and evaluation of culturable bacteria and fungi (including yeast).**

Each replicate was put into a sterile plastic bag with 1L of Phosphate Buffer Saline buffer (8 g/l NaCl, 200 mg/l of KCl, 1.44 mg/l of Na_2_HPO_4_, and 245 mg/l of KH_2_PO_4_, final solution pH=7.4). Then, fruits were bath sonicated (Ultrasonic cleaner, VWR USC100TH; 45 kHz) for 15 min and shaken at room temperature for 20 min at 120 rpm (2 x g). The microbiota washing process was repeated twice to improve the microbiota harvest. Apple fruits outwashes were filtered on a sterile 0.22 µm filters (PALL). Then, to recover microbial cells, filters were put in a 45 mL falcon tube with 30 mL of sterile NaCl solution (8.5 g/L) and washed thoroughly by vortexing (2 min). Aliquots of the microbiota were made from the recovered cells with 30% of glycerol, immediately flash frozen in liquid nitrogen and conserved at -80 °C for biobanking.

Before starting any experiment, the glycerol was eliminated by vacuum filtration (Bio Rad, 900 cu ft/min) of aliquots on a sterile 0.22 µm filters. Before removing filters from the vacuum pump, 5 mL of sterile NaCl (0.85%) solution was poured on the filters to rinse the remaining glycerol. This rinsing process was repeated twice. One aliquot of each sample was 10-fold serially diluted and plated in triplicates on Potato Dextrose Agar (39g/l, Merck, Rhisnes, Belgium) with 100 mg/l of chloramphenicol and Reasoner 2Agar media (Sigma) to respectively estimate culturable fungal and bacterial populations. Colony Forming Units were counted after 7 days of incubation (23°C, 16/8 photoperiod) and used as a proxy of cell concentration to normalise the sample concentrations during bioassays.

**Supplementary material 13a : Taxonomic analyses of harvested microbiota by HTS**

Laboratory Protocol

Two biological replicates of each sample (18 microbiota x 2 replicates) were centrifuged for 10 min at 10,000 x g and the pellet was used as starting material for DNA extraction with FASTDNA spin kit (MP Biomedicals) according to manufacturer’s instructions. The v3-v4 16 rRNA gene and tThe ITS1 region were targeted using respectively Bakt_341F/ Bakt_805R [23] and ITS1-F_KYO2F/ ITS2_KYO2R [24] modified primers with Illumina™ adapters. PCR mix and thermal cycle were identical to previously published protocols [21]. The amplicon presence was checked on 1% gel agarose (90 V, 40 min). Individual tags were added to the PCR products before the sequencing on Illumina MiSeq (2 x 300 nt) according to standard protocols.

Bioinformatics analyses

Quality filtering, denoising, merging, chimera removal, and Amplicon Sequence Variants (ASVs) feature tables were done with DADA2 [26] implemented in q2 with default setting for 16s rRNA. For fungi, forward and reverse read lengths were truncated at positions 232 and 182, respectively, to keep high-quality sequences. Sequences were aligned in q2-MAFFT algorithm [27]; then aligned sequences were used to build phylogenetic trees with FastTree 2 [28], which was further rooted (q2 midpoint-root). Rarefaction curves were computed in q2 diversity alpha-rarefaction with 20 steps at 19,200 and 5,789 maximum sequence depth for 16S rRNA gene and for ITS1 ASVs, respectively. ASVs were taxonomically assigned with vectorized search method VSEARCH [29] in the q2 feature‐classifier plugin. SILVA_132 and UNITE version 8, both at 99% of sequence similarity, were the taxonomic reference databases for bacteria and fungi, respectively. Cytoplasmic contamination sequences (chloroplast and mitochondria) were removed from the bacterial features table with the q2 script taxa filter‐table. The q2‐diversity core‐metrics‐phylogenetic plug‐in was used to calculate alpha (Observed OTUs, Faith Phylogenetic Diversity (Faith PD), Shannon and Pielou’s Evenness) and beta (Bray Curtis, Weighted and Unweighted Unifrac distance metrics) diversity indices on rarefied feature tables using default parameters.

**Supplementary material 13b : Taxonomic analysis of isolated strains**

One millilitre of each suspension was heated during 30s in microwave (750 W) and used directly as DNA templates with ITS1-F [24] and ITS4 [35] primers, following the published protocols. Amplicons were sent to Macrogen (Amsterdam-Zuidoost, Netherland) for purification and dideoxy sequencing (Sanger). BLASTn, analysis against the NCBI ITS databases was carried out to determine the taxonomy of the isolates. The taxonomic assignments were further confirmed in the online BLAST tool UNITE database.

**Supplementary Material 14:** **High resolution graph of Figure 2**

Clustered heatmap of correlations between biological efficacy and taxa abundance in the studied microbiota [applied alone at 10^5^ CFU/ml or co-applied at 3 concentrations (10^4^ CFU/ml, 5.10^4^ CFU/ml and 10^5^ CFU/ml ) with strain K]. Each color represents a correlation, and the strength of the correlation is shown at right side in the legend. Each column represents microbial assigned taxa and each line represents the efficacy of biocontrol assay with the four tested conditions for microbiotas. Blue and red colors indicate positive and negative correlations, respectively.

**Supplementary Material 15:** **High resolution graph of Figure 3**

Clustered heatmap of correlations between biological efficacy of microbiotas [applied alone at 10^5^ CFU/ml or co-applied at 3 concentrations (10^4^ CFU/ml, 5.10^4^ CFU/ml and 10^5^ CFU/ml ) with strain K] and the efficacy of carbon source metabolization for the studied microbiota. Each column represents a carbon source and each line represents the efficacy of biocontrol assay with one tested condition for the microbiotas. Each color represents a correlation, and the strength of the correlation is shown at right side in the legend. Blue and red colors indicate positive and negative correlations, respectively.
